# Anatomically Guided Deep Learning Reconstruction of Accelerated Snapshot CEST MRI

**DOI:** 10.64898/2026.09.28.755205

**Authors:** Emmanuel Akwasi Mensah, Abrar Faiyaz, Giovanni Schifitto, Md Nasir Uddin

**Affiliations:** Department of Biomedical Engineering, University of Rochester, Rochester, NY 14627, USA; Department of Neurology, University of Rochester, Rochester, NY 14642, USA; Department of Electrical & Computer Engineering, University of Rochester, Rochester, NY; Department of Imaging Sciences, University of Rochester, Rochester, NY 14642, USA

**Keywords:** MRI, brain, CEST, APTw, deep learning, compressed sensing

## Abstract

**Purpose:** To determine whether structural MRI information can improve reconstruction of highly accelerated 3D snapshot chemical exchange saturation transfer (CEST) MRI.

**Methods:** Fully sampled brain CEST data were retrospectively undersampled at acceleration factors (AFs) of 4, 6, 8, 10, and 12. We compared reconstruction without structural information with population-level pretraining using T1-weighted (T1w), T2-weighted (T2w), or combined T1w–T2w images, as well as direct conditioning using a co-registered subject-specific T1w image. Performance was evaluated in two held-out healthy subjects using peak signal-to-noise ratio (PSNR), structural similarity index (SSIM), and mean absolute error (MAE) across frequency offsets. Z-spectrum fidelity and derived magnetization transfer ratio asymmetry (MTR_asym_) maps were additionally evaluated within three-dimensional gray- and white-matter masks.

**Results:** Subject-specific T1w conditioning consistently achieved the highest PSNR and SSIM and the lowest MAE across AFs, with greater improvements at higher acceleration. It also improved Z-spectrum agreement, particularly in white matter and combined gray- and white-matter regions across most AFs. Improvements in MTR_asym_ were more modest and varied across tissue types and acceleration factors.

**Conclusion:** Incorporating subject-specific structural information can improve reconstruction of highly accelerated snapshot CEST MRI, particularly at higher acceleration factors. However, improvements in reconstructed source images did not consistently translate into better preservation of derived CEST contrast, highlighting the need for reconstruction strategies that more directly incorporate CEST spectral information.

## 1. Introduction

Chemical exchange saturation transfer (CEST) is an emerging molecular MRI technique that enables indirect sensitivity to endogenous metabolites and exogenous contrast agents containing exchangeable protons [1]. CEST MRI has shown promise for detecting metabolic alterations associated with tumors and neurological disorders, including Alzheimer’s disease, multiple sclerosis, and stroke [2-4]. CEST contrast is generated by applying a radiofrequency saturation pulse at the resonance frequency of an exchangeable solute proton pool, resulting in a reduction in the measured water signal [1]. The magnitude of this signal reduction depends on several factors, including solute concentration, pH, proton-exchange rates, and the applied saturation parameters [5, 6].

A typical CEST acquisition acquires images at multiple frequency offsets, enabling assessment of contrasts involving amide, amine, and hydroxyl protons, as well as nuclear Overhauser enhancement (NOE) [5, 7]. However, the measured CEST signal is influenced by direct water saturation, semisolid magnetization transfer (MT), and magnetic-field inhomogeneity. Acquiring data over a broad range of frequency offsets enables spectral fitting, correction, and quantitative analysis of these effects [8, 9]. Consequently, conventional CEST acquisitions require many frequency-offset images, resulting in long scan times that limit clinical translation [10].

Several approaches have been proposed to accelerate CEST MRI through pulse sequence development and advanced reconstruction methods [11]. One notable example is snapshot CEST, which uses a rapid gradient-echo readout to acquire k-space data for a given frequency offset following a single CEST preparation [12, 13]. However, the evolution of the prepared CEST contrast constrains readout duration and k-space sampling. Further acceleration through k-space undersampling can reduce signal-to-noise ratio (SNR) and introduce undersampling artifacts. Beyond sequence development, conventional compressed sensing methods use various handcrafted priors but may become less effective at higher acceleration factors (AFs) [11, 14, 15]. More recently, deep learning approaches, including image-to-image mapping, unrolled reconstruction networks, and methods that exploit redundancies across the Z-spectrum, have shown promising results for CEST reconstruction [16-18].

In image reconstruction, effective priors can enable the recovery of high-quality images [19, 20]. Nevertheless, few studies have explored the use of commonly acquired MRI contrasts, such as T1-weighted (T1w) and T2-weighted (T2w) images, to facilitate CEST reconstruction. These anatomical images contain structural information that may constrain reconstruction and improve image quality. Multi-contrast reconstruction has previously been used to reconstruct one image contrast using information from another, such as using T1w images to assist T2w image reconstruction [21, 22]. In CEST MRI, Wang et al. incorporated a T2w image into a recurrent feature-sharing network, improving reconstruction of two amide proton transfer offsets [23]. More recently, structural T1w and T2w images have been used to pretrain a super-resolution network, with the learned weights subsequently transferred to CEST reconstruction [24]. However, the potential value of structural information for reconstructing the full CEST Z-spectrum, especially at high acceleration factors, remains less well explored.

The goal of this work was to investigate whether structural MRI could facilitate reconstruction of the full Z-spectrum from accelerated snapshot CEST data. We evaluated two strategies for incorporating structural information. The first used population-level structural pretraining, whereas the second used direct subject-specific conditioning, in which a co-registered T1w image from the same subject was provided as an additional input during CEST reconstruction. We compared these approaches with reconstruction without structural information across multiple AFs.

## 2. Methods

### 2.1 Subjects and Data Acquisition

Fully sampled 3D CEST datasets were acquired from 16 healthy participants using a 3.0 T MAGNETOM Prisma MRI scanner (Siemens Healthineers, Erlangen, Germany) equipped with a 64-channel head coil. Data were acquired using the snapshot CEST sequence [12]. The study was approved by the Institutional Review Board of the University of Rochester, and written informed consent was obtained from each participant before imaging.

The CEST acquisition comprised one unsaturated reference image at -300 ppm and 29 saturation offsets spanning -6.00 to +6.00 ppm, with denser sampling around ±3.5 ppm. The CEST preparation consisted of 36 Gaussian saturation pulses (B_1_ =2.0µ T; pulse duration = 99.84 ms; interpulse delay = 100 ms), followed by a 5000 ms recovery period. The readout parameters were repetition time/echo time (TR/TE) = 4/2 ms, flip angle = 6^∘^, field of view (FOV) = 220 × 220 × 124 mm^3^, spatial resolution = 2 × 2 × 2 mm^3^. Parallel imaging was not used. The total CEST acquisition time was 18 min.

For each participant, a T1w image was acquired using a magnetization-prepared rapid gradient-echo (MPRAGE) sequence [25]. The acquisition parameters were TR/TE = 1840/2.34 ms, FOV = 256 × 256 × 192 mm^3^, and spatial resolution = 1 × 1 × 1 mm^3^.

### 2.2 Retrospective Undersampling and Data Preprocessing

Fully sampled snapshot CEST k-space data were retrospectively undersampled using Cartesian variable-density Poisson-disc (VDPD) sampling at acceleration factors (AFs) of 4, 6, 8, 10, and 12 using the Berkeley Advanced Reconstruction Toolbox (BART) [26]. Coil sensitivity maps were estimated from the fully sampled data using ESPIRiT in BART [27]. The undersampled multicoil k-space was combined using the estimated sensitivity maps to produce the aliased complex images for network input. The sampling masks used are shown in Supporting Information Figure S1, and representative undersampled CEST source-images are presented in Supporting Information Figure S2.

For subject-specific structural conditioning, each subject’s T1w image was registered to CEST space using the fully sampled data as the reference. A single transformation was estimated and applied across all AFs. Registration was performed using FMRIB’s Linear Image Registration Tool (FLIRT) in FSL [28]. Two subjects were excluded due to poor registration that persisted despite tuning of the registration parameters. A representative registration is provided in Supporting Information Figure S4. Following coregistration, the T1w image was normalized by its 95th-percentile intensity.

### 2.3 Deep Learning Reconstruction Architecture

CEST images were reconstructed using a three-dimensional denoising convolutional neural network (3D DnCNN) [29]. Each complex-valued CEST image was represented as real and imaginary channels with the co-registered T1w image concatenated as a third channel for subject-specific conditioning. All other configurations received only the two CEST channels. The network generated two output channels corresponding to the real and imaginary components. They were combined to form the magnitude image.

The network comprised 12 convolutional layers operating at the original spatial resolution. The first layer used a 3 × 3 × 3 convolution to map the input channels to 64 feature maps, followed by a rectified linear unit (ReLU) activation. Each of the subsequent 10 layers applied a 3 × 3 × 3 convolution producing 64 feature maps, followed by batch normalization and ReLU activation. The final layer used a 3 × 3 × 3 convolution to map the 64 feature maps to two output channels. All convolutional layers used a stride of one and padding of one, preserving the spatial dimensions throughout the network. The network was trained using residual learning. Figure 1 shows the network architecture.

**Figure 1.**
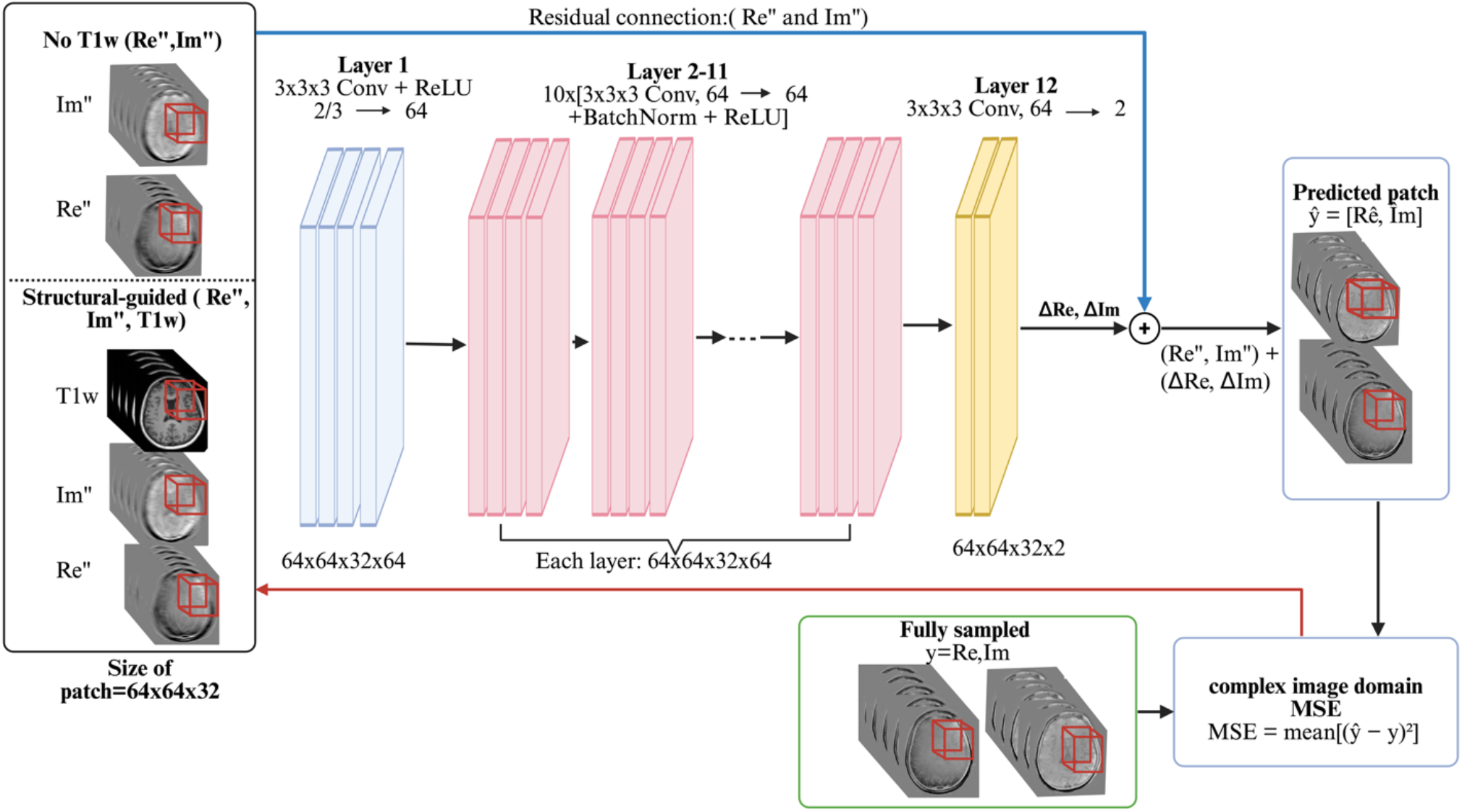
Architecture of a 3D DnCNN operating on complex-valued data. In the anatomically guided configuration, the co-registered T1w image is supplied as an additional third input channel. The first convolutional layer maps the input channels to 64 feature maps, followed by 10 intermediate layers, each applying a 3×3×3 convolution, batch normalization, and a ReLU activation, and a final convolutional layer mapping back to 2 channels. The network is trained with residual learning. The baseline configuration is identical but omits the T1w input channel.

### 2.4 Model Training

The 14-subject dataset was divided into nine for training, three for validation, and two for testing. Across five AFs, this yielded 45 training, 15 validation, and 10 test volumes per frequency offset. Given the limited dataset, we used patch-based training. From each volume, we randomly extracted 200 3D patches of size 64 × 64 × 32 voxels. We trained separate models for each frequency offset, first training on the unsaturated offset for 60 epochs, then using its weights to initialize training on the remaining models for up to 30 epochs. This transfer-learning scheme avoided training each offset from scratch and exploited the shared low-level image features across offsets. We employed a cosine learning-rate schedule with an eight-epoch linear warm-up, using a peak learning rate of 3 × 10^−4^ for the unsaturated offset and 1 × 10^−4^ during transfer learning. The network was optimized with AdamW optimizer using mean squared error (MSE) loss. Training and validation MSE curves for the unsaturated offset are presented in Supporting Information Figure S3.

The models were implemented in PyTorch 2.9.0 and trained on an Ubuntu workstation equipped with an Intel Core i9-10920X CPU, 125 GB of system memory, and an NVIDIA RTX A6000 GPU.

### 2.5 Structural pretraining

For structural pretraining, we obtained 200 T1w and 200 T2w images from the ADNI database [30, 31]. First, we Fourier-transformed each image, retrospectively undersampled k-space using dimension-matched VDPD masks and applied the inverse transform to obtain complex aliased images. Given the larger dataset, 50 patches of size 64 × 64 × 32 voxels were extracted from each volume. Using an 80/20 training–validation split, three two-channel DnCNN models were pretrained for 60 epochs with a batch size of 32 on T1w, T2w, or combined T1w–T2w data.

We then used each pretrained model to initialize CEST training, first on the unsaturated offset and subsequently, by weight transfer, on the remaining offsets. During fine-tuning for the unsaturated offset, we evaluated peak learning rates of 3×10^−4^, 1×10^−4^, 3×10^−5^, and selected the best. All CEST-stage training parameters were held identical to those used for the model with no T1w.

### 2.6 Evaluation

We compared reconstructed images against the fully sampled reference using the peak signal-to-noise ratio (PSNR), structural similarity index (SSIM), and mean absolute error (MAE). We also evaluated the reconstructions in terms of CEST quantification. All reconstructed and reference data underwent post-processing using code adapted from the CEST-sources (https://github.com/cest-sources/CEST_EVAL). For each voxel, a local *B*_0_ offset was estimated from the minimum of a smoothing-spline-fitted Z-spectrum, followed by frequency-shifting, linear interpolation, and normalization by the unsaturated reference image [32]. We then derived magnetization transfer ratio asymmetry (MTR_asym_) maps by using [33]:

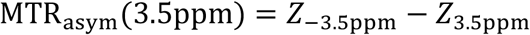

where *Z*_−3.5ppm_ is the reference image at -3.5ppm, and *Z*_3.5ppm_ is the saturated image at -3.5ppm.

We also compared Z-spectra and MTR_asym_ maps within gray matter (GM) and white matter (WM). GM and WM masks were derived from the T1w image registered to CEST space using FMRIB’s Automated Segmentation Tool (FAST) and thresholded at a tissue probability of 0.8 [34].

## 3. Results

### 3.1 Quantitative and qualitative assessment of CEST source images

Table 1 summarizes PSNR, SSIM, and MAE for the reconstructed CEST source images. Subject-specific T1w conditioning yielded the highest PSNR and SSIM and the lowest MAE across all AFs. At AF12, subject-specific T1w conditioning achieved a PSNR of 34.01 ± 0.97 dB and SSIM of 0.9145 ± 0.0147, compared with 33.56 ± 0.73 dB and 0.9006 ± 0.0078, respectively, without structural information. MAE was also lower with subject-specific T1w conditioning (0.0140 ± 0.0016) than without structural information (0.0150 ± 0.0012). In contrast, population-level structural pretraining produced relatively small and inconsistent differences relative to reconstruction without structural information. Frequency-resolved metrics were calculated within each subject’s 3D WM masks and averaged across the two test subjects showed similar findings (Supporting Information Figure S4). Across offsets, subject-specific T1w conditioning achieved higher PSNR and SSIM and lower MAE than the other methods, including at higher AFs.

**Table 1.** Quantitative assessment using Peak signal-to-noise ratio (PSNR; dB), structural similarity index measure (SSIM), and (C) mean absolute error (MAE) of reconstructed CEST source images for a selected slice across all frequency offset. For each subject and reconstruction method, each metric was calculated at every frequency offset and then averaged across offsets. Values are reported as the mean ± standard deviation across two test subjects (n = 2). Bold values indicate the best-performing method at each acceleration factor (AF). Higher PSNR and SSIM values and lower MAE values indicate better reconstruction quality.

|  | Method | AF4 | AF6 | AF8 | AF10 | AF12 |
| --- | --- | --- | --- | --- | --- | --- |
| PSNR | No T1w | 37.47 ± 1.19 | 35.71 ± 0.87 | 34.82 ± 0.78 | 34.39 ± 0.91 | 33.56 ± 0.73 |
| | T1w pretraining | 37.55 ± 1.09 | 35.88 ± 0.87 | 34.79 ± 0.75 | 34.24 ± 0.84 | 33.53 ± 0.82 |
| | T2w pretraining | 37.42 ± 0.73 | 35.70 ± 0.73 | 34.62 ± 0.66 | 34.11 ± 0.85 | 33.37 ± 0.67 |
| | T1w+T2w pretraining | 37.43 ± 0.85 | 35.77 ± 0.64 | 34.70 ± 0.72 | 34.30 ± 0.85 | 33.62 ± 0.62 |
|  | Subject-specific T1w | **37.64 ± 1.15** | **35.92 ± 1.05** | **35.10 ± 0.87** | **34.57 ± 1.00** | **34.01 ± 0.97** |
| SSIM | No T1w | 0.9546 ± 0.0067 | 0.9341 ± 0.0073 | 0.9185 ± 0.0080 | 0.9145 ± 0.0112 | 0.9006 ± 0.0078 |
| | T1w pretraining | 0.9547 ± 0.0063 | 0.9360 ± 0.0061 | 0.9199 ± 0.0059 | 0.9141 ± 0.0074 | 0.9014 ± 0.0077 |
| | T2w pretraining | 0.9542 ± 0.0038 | 0.9343 ± 0.0067 | 0.9171 ± 0.0081 | 0.9109 ± 0.0113 | 0.8981 ± 0.0085 |
| | T1w+T2w pretraining | 0.9541 ± 0.0057 | 0.9350 ± 0.0051 | 0.9182 ± 0.0079 | 0.9140 ± 0.0099 | 0.9018 ± 0.0069 |
|  | Subject-specific T1w | **0.9586 ± 0.0070** | **0.9414 ± 0.0105** | **0.9288 ± 0.0116** | **0.9236 ± 0.0146** | **0.9145 ± 0.0147** |
| MAE | No T1w | 0.0096 ± 0.0012 | 0.0118 ± 0.0012 | 0.0130 ± 0.0012 | 0.0136 ± 0.0014 | 0.0150 ± 0.0012 |
| | T1w pretraining | 0.0096 ± 0.0012 | 0.0116 ± 0.0011 | 0.0130 ± 0.0011 | 0.0137 ± 0.0013 | 0.0150 ± 0.0013 |
| | T2w pretraining | 0.0097 ± 0.0008 | 0.0118 ± 0.0010 | 0.0132 ± 0.0011 | 0.0140 ± 0.0015 | 0.0153 ± 0.0011 |
| | T1w+T2w pretraining | 0.0096 ± 0.0010 | 0.0117 ± 0.0009 | 0.0132 ± 0.0011 | 0.0137 ± 0.0013 | 0.0149 ± 0.0010 |
|  | Subject-specific T1w | **0.0093 ± 0.0012** | **0.0114 ± 0.0014** | **0.0125 ± 0.0014** | **0.0131 ± 0.0016** | **0.0140 ± 0.0016** |

Figure 2A shows reconstructed CEST source images at 3.5 ppm for a representative test subject. At AF4 and AF6, all methods produced comparable image quality. At AF8–AF12, the subject-specific T1w conditioning better preserved anatomical features, including ventricular boundaries and surrounding tissue structures. The corresponding absolute-error maps showed generally lower errors with subject-specific T1w conditioning (Figure 2B).

**Figure 2.**
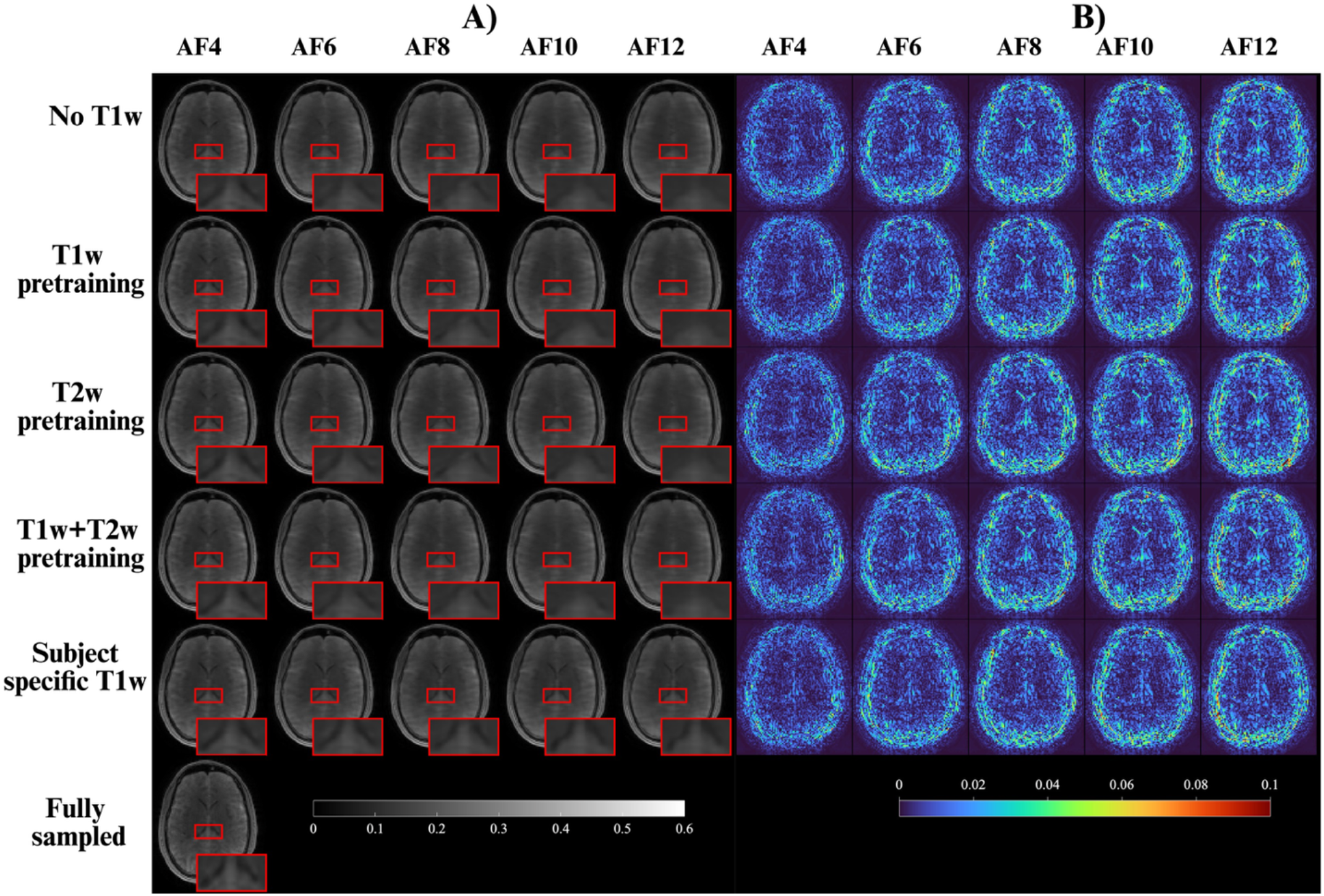
Qualitative comparison of CEST source-image reconstruction methods across acceleration factors. (A) Reconstructed source images from the same representative axial slice at the +3.5-ppm are shown for AFs of 4, 6, 8, 10, and 12. Rows correspond to reconstruction with no T1w, T1w pretraining, T2w pretraining, T1w+T2w pretraining, and reconstruction using the subject-specific T1w image, followed by the fully sampled data. Red boxes indicate the regions displayed in the magnified insets. (B) Corresponding voxel wise absolute-error maps, calculated as the absolute difference between each reconstructed image and the fully sampled reference. Lower error values indicate greater agreement with the fully sampled image. T1w, T1-weighted; T2w, T2-weighted; AF, acceleration factor.

### 3.2 Full Z-spectrum and CEST maps comparison

In WM, subject-specific T1w conditioning yielded the lowest Z-spectrum MAE across all evaluated AFs (Figure 3A). In GM, it produced comparable or lower MAE at AF4–AF10, but a higher MAE at AF12 than the other methods, except for T1w pretraining (Figure 3B). Within the combined GM+WM mask, it produced comparable MAE at AF4 and the lowest MAE at higher AFs (Figure 3C). These results indicate improved overall Z-spectrum fidelity with subject-specific T1w conditioning.

**Figure 3.**
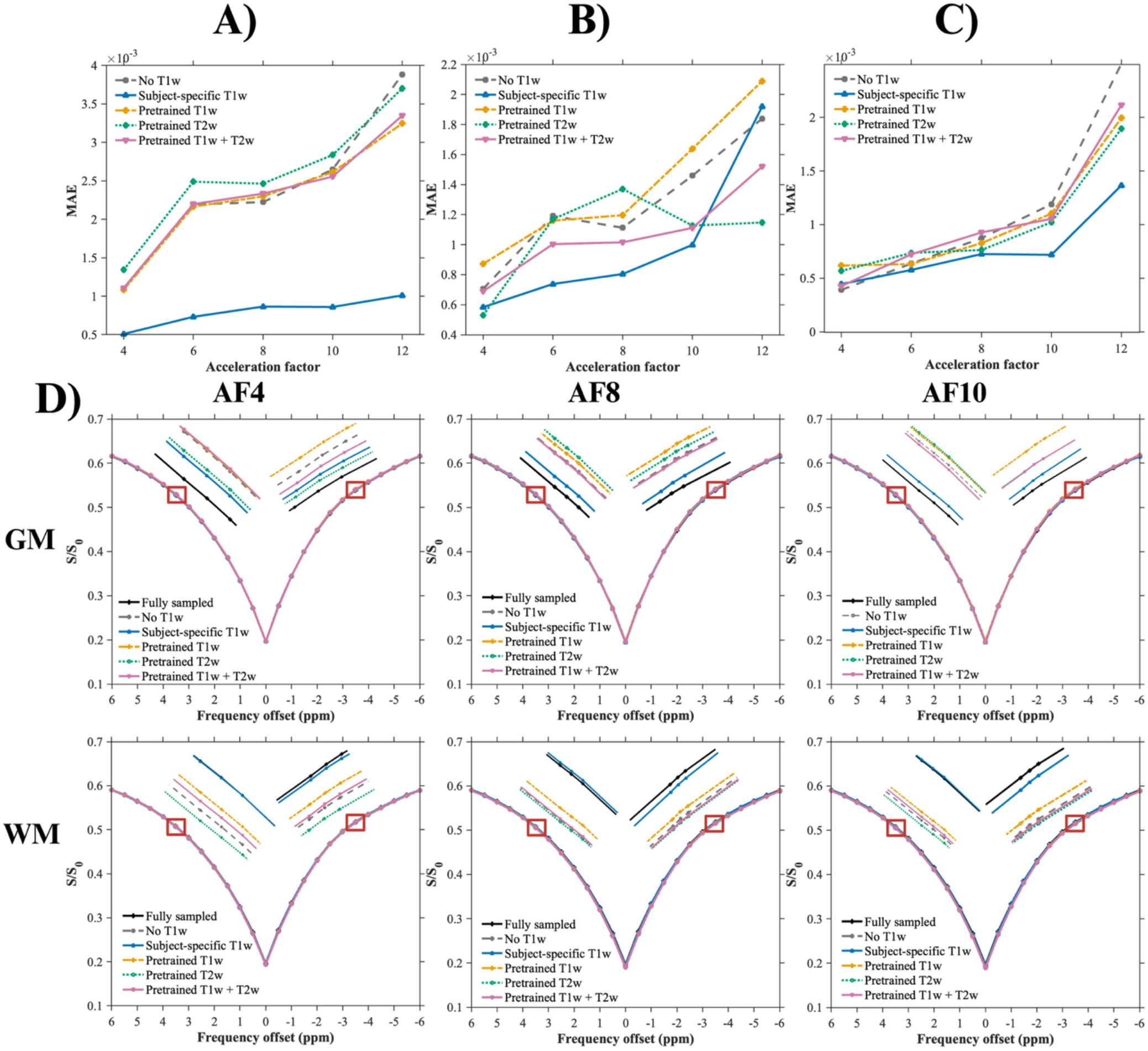
Tissue-specific assessment of reconstructed CEST Z-spectrum fidelity across acceleration factors. (A–C) Mean absolute error (MAE) between the reconstructed and fully sampled Z-spectra at selected acceleration factors (AF), calculated using all voxels within the (A) white-matter (WM), (B) gray-matter (GM), and (C) combined GM+WM masks. Values represent the mean across two test subjects; lower MAE indicates greater agreement with the fully sampled reference. (D) Mean Z-spectra within the three-dimensional GM (top row) and WM (bottom row) masks for a representative test subject at AF4, AF8, and AF10. Curves compare the fully sampled reference with reconstruction with no T1w, subject-specific T1w, T1w pretraining, T2w pretraining, and combined T1w and T2w pretraining.

Figure 3D compares the mean GM and WM Z-spectra for a representative test subject at AF4, AF8, and AF10. Subject-specific T1w conditioning showed greater consistency with the fully sampled spectra in both GM and WM, whereas the other methods generally overestimated the GM Z-spectrum and produced relatively lower WM Z-values.

Figure 4 evaluates whether these differences propagated to the derived *MTR*_asym_ maps. The reconstructed maps showed comparable spatial distributions across methods and AFs (Figure 4A), while the corresponding absolute-error maps generally showed lower errors with subject-specific T1w conditioning at AF6–AF12 (Figure 4B).

**Figure 4.**
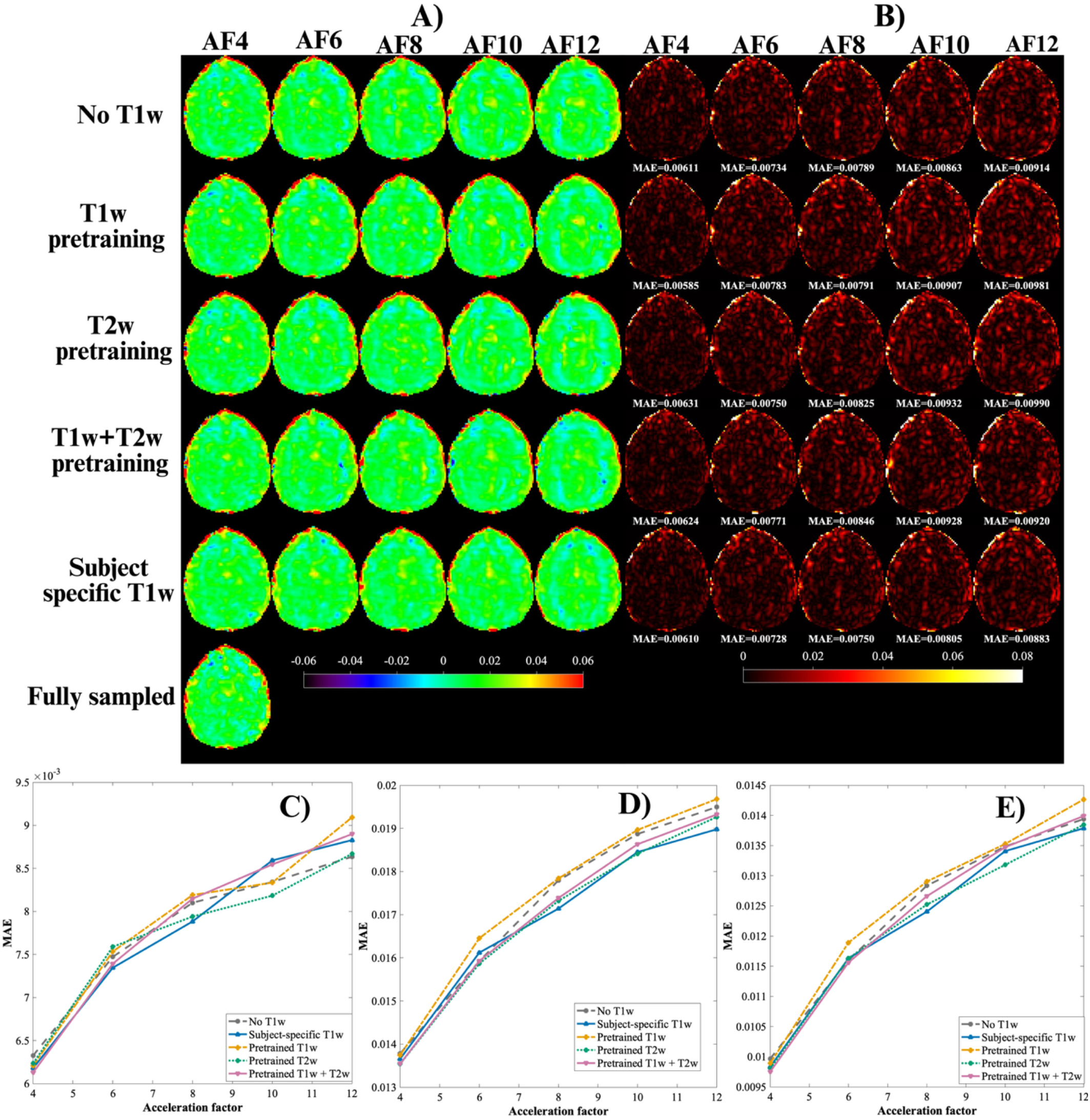
Qualitative and quantitative assessment of MTRasym across acceleration factors for a test subject. *(A) MTRaysm (*3.5 ppm) maps from a representative axial slice at acceleration factors (AFs) of 4, 6, 8, 10, and 12. Rows correspond to reconstruction with no T1w, T1w pretraining, T2w pretraining, combined T1w and T2w pretraining, subject-specific T1w, and the fully sampled data. (B) Corresponding voxelwise absolute-error maps relative to the fully sampled MTRasym *map; (C–E) MTRaysm MAE at various acceleration factors within the (C) white-matter, (D) gray-matter, and (E) combined gray- and white-matter masks. Lower MAE indicates greater agreement with the fully sampled reference*.

The tissue-specific *MTR*_asym_ MAE analysis showed that the effect of the subject-specific T1w input was not uniform across masks and AFs, although modest improvements were observed in several conditions. In WM, the subject-specific T1w conditioning yielded the lowest MAE at AF4, AF6, and AF8, with smaller differences at AF10 and AF12 (Figure 4C). In GM, errors were comparable at AF4 and AF6, whereas subject-specific T1w conditioning yielded lower MAE at AF8–AF12 (Figure 4D). Within the combined GM+WM mask, modest reductions were observed at several AFs (Figure 5E). Overall, subject-specific T1w conditioning improved source-image and Z-spectrum fidelity consistently, while improvements in derived *MTR*_asym_ were smaller and dependent on tissue type and AF.

## 4. Discussion

In this study, we investigated whether structural MRI information could improve reconstruction of accelerated 3D snapshot CEST data. We compared population-level structural pretraining with direct conditioning using a co-registered subject-specific T1w image. Across AFs from 4 to 12, subject-specific T1w conditioning consistently improved source-image reconstruction, with larger differences at higher AFs. It also improved Z-spectrum agreement, particularly in WM and the combined GM+WM mask. However, improvements in *MTR*_asym_ were smaller and varied across tissues and AFs. These findings suggest that subject-specific anatomical information can improve highly undersampled CEST reconstruction, although improved source-image reconstruction does not necessarily translate into comparable improvement in derived CEST contrast.

### 4.1 Impact of subject-specific T1w information on CEST image reconstruction

The consistent improvement with subject-specific T1w conditioning suggests that how structural information is incorporated into the reconstruction matters. Pretraining transfers population-level structural features through the initial network weights, which may be altered during subsequent CEST fine-tuning. In contrast, direct conditioning provides access to the subject’s anatomy throughout reconstruction, which may become increasingly valuable at higher AFs for recovering missing spatial information and preserving tissue boundaries.

However, the Z-spectrum and MTR_asym_ analyses showed that the benefit was not uniform across tissues and AFs. One possible explanation for the reduced performance in GM at AF12 is that, under severe undersampling, structural information influenced the reconstruction more strongly than the CEST signal. Spatial-feature weighting or CEST-specific spectral constraints may therefore help preserve anatomical detail without compromising CEST-specific information.

The smaller improvement in *MTR*_asym_ may also reflect how this measure is calculated. *MTR*_asym_ depends on the difference between signals acquired at frequency offsets on opposite sides of the water resonance. Therefore, small reconstruction errors at either offset can propagate into the resulting *MTR*_asym_ map.

### 4.2 Relationship to previous work

CEST data are inherently SNR-limited because saturation reduces the measured water signal. Further undersampling reduces acquisition time but increases reconstruction difficulty. A co-registered T1w image can constrain this reconstruction at high AFs by providing subject-specific tissue boundaries and anatomical structure. Although T1w imaging requires a separate acquisition, it is commonly included in brain MRI protocols [35, 36]. Existing CEST reconstruction methods have primarily exploited redundancies across saturation offsets [16, 18, 37-39], while using one MRI contrast to assist reconstruction of another is established in multi-contrast MRI [21, 40, 41].

Building on prior CEST studies [23, 24], we compared direct subject-specific T1w conditioning with population-level pretraining. The results suggest that direct structural conditioning may be more useful when a co-registered T1w image is available. However, subject-specific T2w and combined T1w–T2w conditioning were not evaluated. Future studies could compare these structural inputs to identify the most effective prior. The relatively small benefit of structural pretraining should also be interpreted within the pretraining data and network architecture used here.

### 4.3 Limitations and future directions

This study has several limitations. First, reconstruction performance was evaluated in only two healthy test subjects, although both were completely held out from training and validation. Larger and multicenter cohorts are therefore required to establish the reproducibility and generalizability of the findings.

Second, the study used retrospective undersampling of fully sampled CEST data. Future studies should implement prospective undersampling to determine whether subject-specific T1w conditioning improves reconstruction of accelerated-acquisition CEST. Validation in patients with neurological disorders such as brain tumors is also needed to determine whether structural priors preserve disease-related CEST abnormalities.

Finally, acquiring T1w and CEST images with matched geometry, including FOV, orientation, and matrix size, could reduce the registration errors and improve spatial correspondence. Modality dropout and learned feature-weighting mechanisms could further reduce sensitivity to misregistration and prevent excessive reliance on structural information.

## 5. Conclusion

This study provides an initial demonstration that direct incorporation of a co-registered subject-specific T1w image can improve three-dimensional snapshot CEST reconstruction. Subject-specific T1w conditioning outperformed reconstruction without structural information and structural-image pretraining across the evaluated acceleration factors. However, improvements in MTR_asym_ were smaller and varied with tissue type and acceleration factors, indicating that improved source-image reconstruction does not ensure uniform preservation of derived CEST contrast. Larger prospective studies, including patients and data acquired across different scanners and protocols, are needed to confirm the generalizability and robustness of these findings.

## Acknowledgments

This study was supported by a grant from the National Institutes of Health (5DP1DK139798-02).

The authors sincerely thank all study participants and study coordinators for their invaluable contributions to this work. We also gratefully acknowledge Professor Moritz Zaiss for his valuable discussions and insights regarding the snapshot CEST sequence.

## Data Availability Statement

The data that support the findings of this study are available on request from the corresponding author. The data are not publicly available due to privacy or ethical restrictions.

