## Supporting Information for "Anatomically Guided Deep Learning Reconstruction of Accelerated Snapshot CEST MRI"

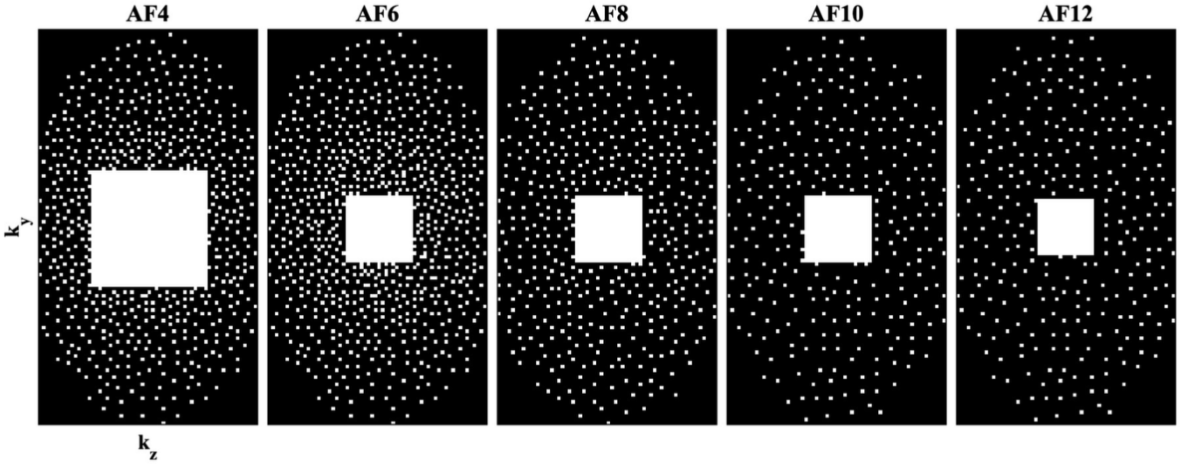


***Supporting Information Figure S1A:*** *Variable-density Poisson-disc sampling masks with elliptical coverage and fully sampled calibration regions in the phase-encoding plane, used for retrospective undersampling at acceleration factors of 4, 6, 8, 10, and 12.*

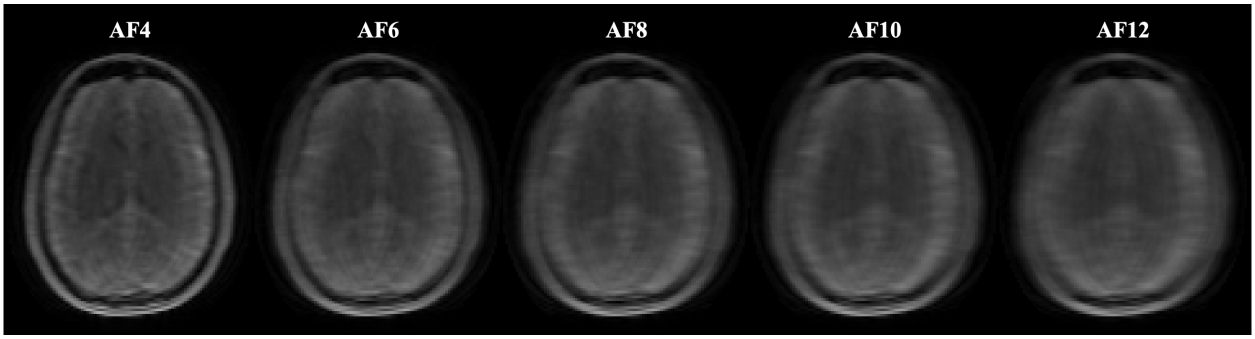


***Supporting Information Figure S1B.*** *Retrospectively undersampled CEST source images at 3.5 ppm from a representative test subject, shown at acceleration factors of 4, 6, 8, 10, and 12, respectively.*

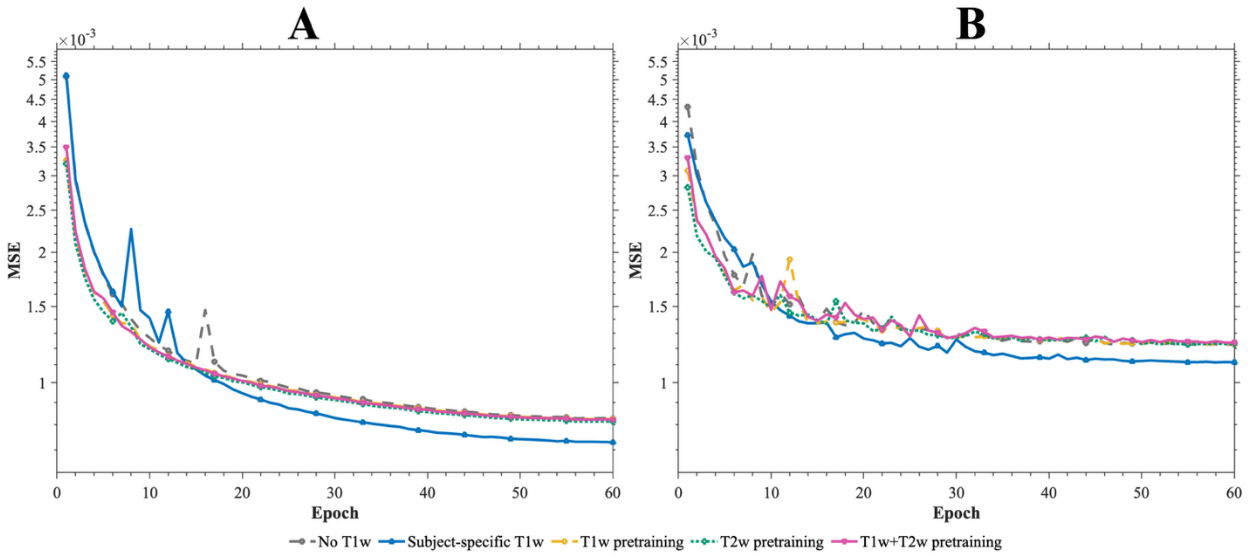


*Supporting Information* ***Figure S2****. Training (A) and validation (B) MSE across 60 epochs for the unsaturated offset; this model was subsequently fine-tuned for the remaining frequency offsets for each method.*

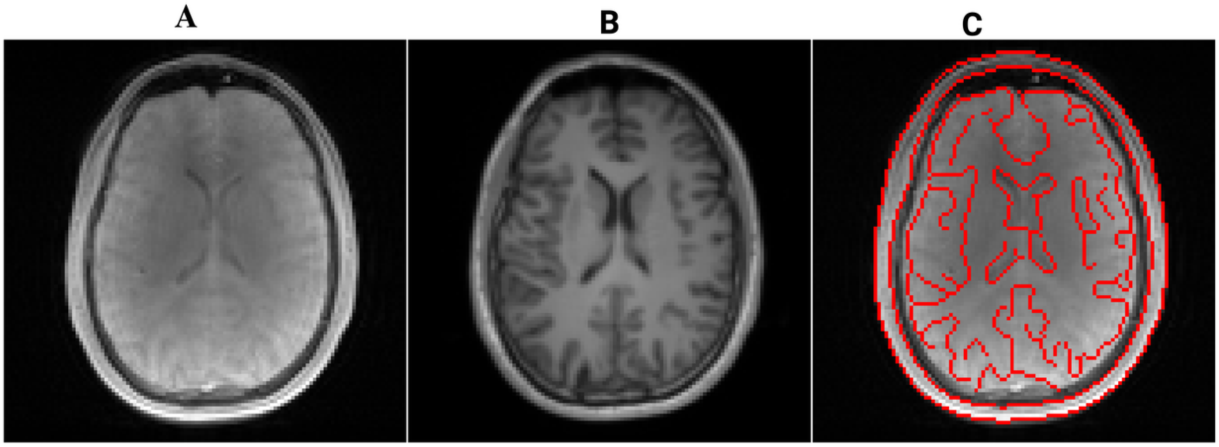


***Supporting Information Figure S3****. Visual assessment of T1w-to-CEST registration for one test subject. (A) Selected axial unsaturated CEST image, (B) corresponding registered T1w image, and (C) T1w-derived anatomical edges overlaid on the CEST image to assess spatial correspondence.*

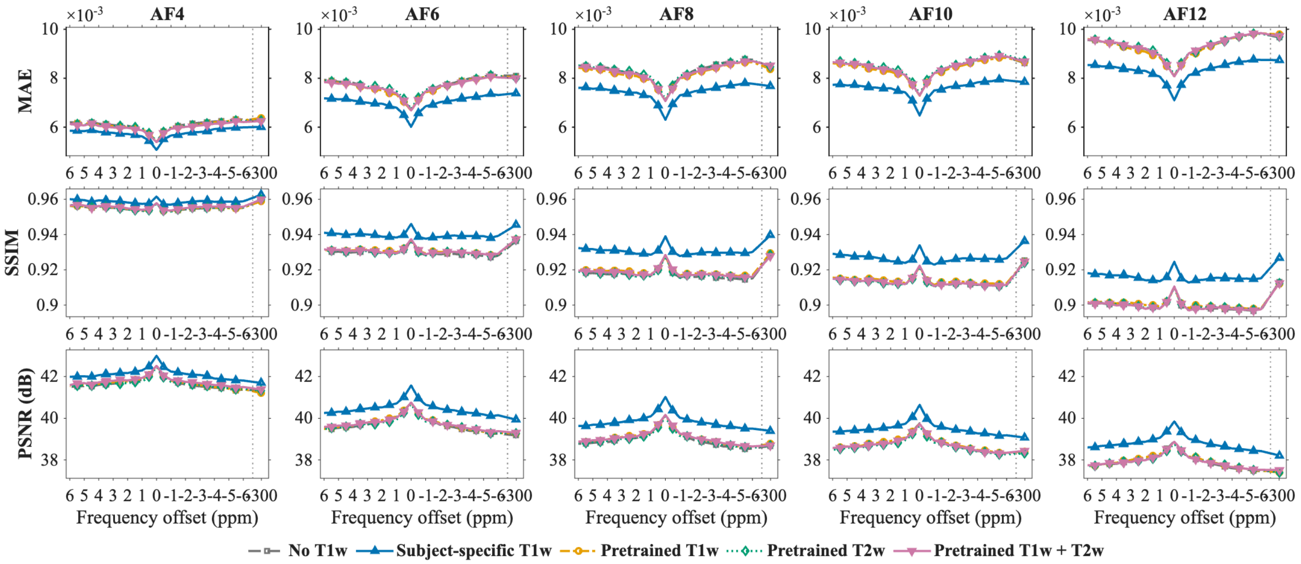


***Supporting Information Figure S4***. *Frequency-resolved quantitative assessment of CEST source-image reconstruction quality within the 3D white-matter (WM) mask for test subjects. Columns correspond to acceleration factors (AFs) of 4, 6, 8, 10, and 12, whereas rows show mean absolute error, structural similarity index measure (SSIM), and peak signal-to-noise ratio (PSNR), respectively.*
